# Development of the “EASY-HSV (Efficient And Simple high-Yield Herpes Simplex virus Vector) system” based on HSV-1 genome-maintaining HEK293 cells

**DOI:** 10.1101/2024.12.05.626320

**Authors:** Fumio Maeda, Shungo Adachi, Tohru Natsume

## Abstract

Herpes simplex virus (HSV) amplicon vector is a promising molecular tool with a large transgene capacity of 150 kb. However, the conventional vector production method is labor-intensive and expensive, limiting its application. Here, we generated HEK293 cells harboring the HSV genome (293/HSV) and developed an EASY-HSV (Efficient And Simple high-Yield Herpes Simplex virus Vector) system in which 293/HSV cells are transfected with only two plasmids.

## Main text

Herpes simplex virus (HSV) amplicon vector, derived from HSV type 1, carries a concatemeric form of DNA plasmid containing transgenes (amplicon plasmid) instead of the viral genome. The HSV amplicon vector has unique advantages for gene delivery, including: (1) a large transgene capacity of up to 150 kb, (2) broad cell tropism, (3) minimal toxicity, inducing only weak adaptive immune responses, and (4) stability and long-term expression ^1 2^. In particular, its large transgene capacity offers a substantial advantage over other viral vectors for gene therapy. However, the conventional vector production method is labor-intensive and expensive, limiting the use of this vector ^1^.

The conventional HSV vector production method involves transfection of the following components: (1) the HSV-1 genome lacking a packaging sequence (*pac*) in viral particles and essential viral gene ICP27; (2) an amplicon plasmid containing the genes of interest (GOI), a *pac* sequence, and the HSV-1 ori sequence (*oriS*); and (3) a plasmid encoding ICP27 (pICP27), into Vero cells expressing the ICP27 gene with its own promoter ^1^. One issue is that the large and fragile HSV-1 genome (>150 kb) is maintained with bacterial artificial chromosome (HSVBAC), which requires purification during each round of vector production using ultracentrifugation and is difficult to introduce into cells through transfection, resulting in low vector yields ^1, 3^. The other issue is that Vero cells have low gene transfection efficiency, requiring expensive transfection reagents ^4^. To address these issues, we selected HEK293 cells with high gene transfection efficiency as the vector-producing cells and established HEK293 cells stably maintaining the HSV-1 genome (293/HSV cells). Using the 293/HSV cells, we developed an efficient, low-cost vector production system, requiring transfection with only an amplicon plasmid and pICP27, designated the “EASY-HSV (Efficient And Simple high-Yield Herpes Simplex virus Vector) system” (Fig. 1a).

**Figure 1.**
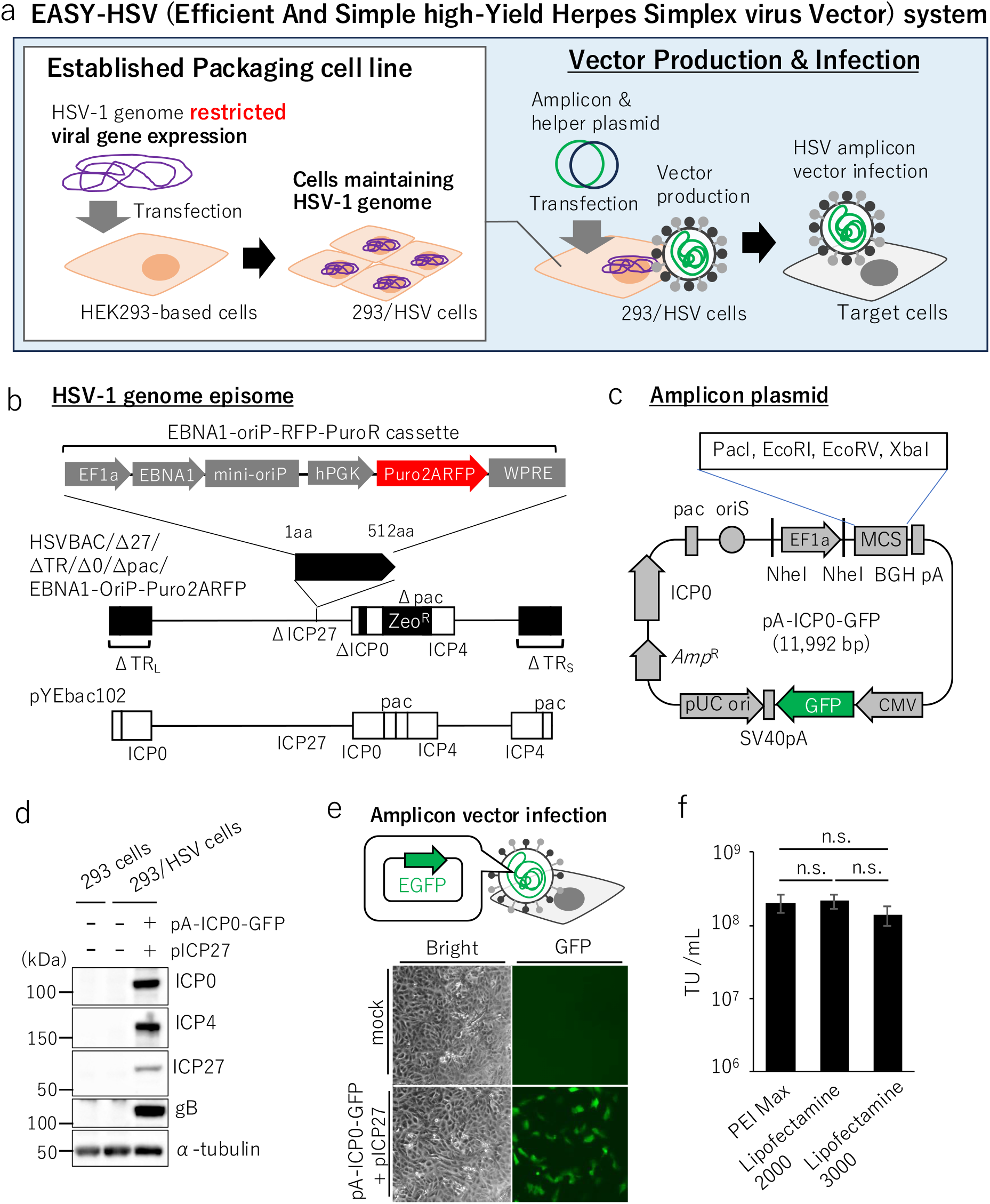
HSV amplicon vector generated using the EASY-HSV system. (a) HSV amplicon vector production methods. (b) Schematic diagram of HSVBAC/Δ27/ΔTR/Δ0/Δpac/EBNA1-oriP-Puro2ARFP (top) constructed from parental HSV-1 BAC pYEbac102 (bottom). The black box indicates the nonsense mutation (ICP0) or deletion (others). TR_L_: terminal region L, TR_S_: terminal region S, *pac*: packaging sequence, Zeo^R^: zeocin-resistant gene. (c) Schematic diagram of the amplicon plasmid constructed in this study. (d) HEK293 and 293/HSV cells were mock-transfected or transfected with pICP27 and pA-ICP0-GFP and analyzed by immunoblot analysis with the indicated antibodies. (e) Fluorescent microscopy images showing Vero cells treated for 18 h with the supernatant purified from mock-transfected cells or cells transfected with pICP27 and pA-ICP0-GFP. (f) Vero cells were infected for 18 h with HSV amplicon vector stocks prepared with the indicated transfection reagent. TU: transduction unit. Data are shown as the means ± standard error of the results of three independent experiments and are expressed relative to the mean determined for Lipofectamine 2000, which was normalized to 1. Statistical analysis was performed by one-way analysis of variance (ANOVA) and Tukey’s test.

First, we constructed a recombinant HSV-1 genome, restricted viral regulatory gene expression to prevent cell death, and enabled autonomous replication in cells to avoid dilution of the viral genome due to cell division. HSV gene expression is a tightly regulated, ordered cascade, which begins with the production of the immediate-early (IE) ^5^. A previous report showed that deletion or mutation of IE genes ICP0, ICP4, ICP22, and/or ICP27 decreased viral gene expression globally in HSV-1-infected cells ^6^. To repress viral gene expression, we deleted ICP27 and the terminal regions, which contain ICP0 and ICP4, and inserted a nonsense mutation into ICP0 in the internal region (Fig. 1b). To avoid dilution of the viral genome due to cell division, we focused on the episomal vector system based on the stable autonomous replication and extrachromosomal persistence of the 172 kb human herpes virus Epstein–Barr virus (EBV) ^7^. The episomal vector system requires the EBV latent origin of replication (oriP) and the EBV nuclear antigen 1 (EBNA-1) protein; EBNA-1 protein connects to repetitive DNA recognition elements within oriP and cellular chromatin, promoting nuclear retention ^7^. We replaced the ICP27 locus in HSVBAC with the gene cassette containing EBNA-1, minimalized oriP (mini-oriP)^7^, the puromycin-resistance gene, and monomeric red fluorescence protein (mRFP) (Fig. 1b). Finally, by deleting a *pac* sequence in the internal region, we constructed HSVBAC/Δ27/ΔTR/0n212/Δpac/EBNA1-oriP-Puro2ARFP (Fig. 1b). HSVBAC was transfected with HEK293 cells and cloned into 293/HSV cells, and mRFP expression was confirmed (Supplementary Fig. 1a). We also constructed an amplicon plasmid (pA-ICP0-GFP) encoding the multi-cloning site (MCS) for the GOI, enhanced green fluorescence protein (EGFP), and ICP0, which inhibits the decrease of amplicon vector-mediated transgene expression ^8^ (Fig. 1c).

Next, we examined 293/HSV cells that produce HSV amplicon vectors. 293/HSV cells were mock-transfected or transfected with pA-ICP0-GFP and pICP27. Immunoblot analysis confirmed that the HSV-1 genes and EGFP were expressed in the transfected cells (Fig. 1d, Supplementary Fig. 1b and c). Vero cells were then treated with the supernatant purified from mock-transfected or transfected cells (Fig. 1e). EGFP expression was detected in cells treated with the supernatant of pA-ICP0-GFP- and pICP27-transfected cells (Fig. 1e). The same level of EGFP expression was also detected with different transfection reagents (Fig. 1f, Table 1). No significant effect of the cultivation period of 293/HSV cells was observed (Supplementary Fig. 1c).

**Table 1:** Transduction unit of vector stocks.

| **Reagent** | **Transduction unit** |
| --- | --- |
| PEI | $2.24 \pm 0.622 \times 10^8$ |
| L2000 | $2.37 \pm 0.505 \times 10^8$ |
| L3000 | $1.53 \pm 0.451 \times 10^8$ |

Our results indicated the successful establishment of HEK293 cells that maintain the HSV genome and confirmed that 293/HSV cells transfected with the amplicon plasmid and pICP27 produce high-titer HSV amplicon vectors (>10^8^ transduction units (TU) from 10^7^ cells) more rapidly and cheaply than the conventional method, which requires purification of the HSV genome ^4^. A limitation of our system is that modifying viral gene products, including viral particle compartments, requires recombination of HSVBAC and the establishment of cells that maintain the recombinant HSV genome.

Our EASY-HSV system uses HEK293 cells as vector production cells and polyethylenimine (PEI) as a transfection reagent. This combination is used to manufacture lentivirus or AAV vectors, which indicates that these manufacturing techniques can also be used to manufacture HSV amplicon vectors, expediting clinical studies using HSV amplicon vectors.

## Acknowledgments

We thank Y. Kawaguchi for providing HSVBAC pYE102bac and Vero cells; G. Smith for providing *Escherichia coli* strain GS1783; Yoshihiko Miwa for providing pEB6CAG; Kosuke Yusa for providing pKLV2-U6gRNA5(BbsI)-PGKpuro2ABFP-W; Nikolaus Osterrieder for providing pEPkan-S; H. Suenaga and K. Shinya for kindly allowing us to use their cell sorter SH800S. We thank Edanz (https://jp.edanz.com/ac) for editing a draft of this manuscript. This study was supported by JSPS KAKENHI Grant Number JP20K20177, JP24K21113 (to F.M.), and 22H02225 (to S.A.); JST CREST Grant Number JPMJCR20E6 (to S.A.); and AMED CREST Grant Number 23gm1610010h0002 (to S.A.).

## Methods

### Cells

Vero cells ^9^ and Flp-In^™^ T-REx^™^ 293 cells (R78007; Thermo Fisher Scientific, Sunnyvale, CA, USA) were maintained in Dulbecco’s modified Eagle’s medium (DMEM; Wako, Osaka, Japan) containing 10% fetal bovine serum (FBS; Biosera, Nuaille, France).

### Antibodies

The following antibodies were used for immunoblotting analysis: mouse monoclonal antibodies against gB (P1105; Virusys, MA, USA), ICP0 (#56985; Santa Cruz Biotechnology, CA, USA), ICP4 (ab6514; Abcam, Cambridge, UK), ICP27 ([H1113], ab53480; Abcam), and α-tubulin (#4970; #T6199-200UL; Sigma-Aldrich, MO, USA).

### Plasmids

pICP27 was constructed by retrieving the fragment that contains the ICP27 promoter and the coding region between the *Bam*HI–*Sac*I sites (GenBank: GU734771.1 map positions 113,213–115,636) ^10^ from pYEbac102 (HSV-1 BAC) ^9^ and inserting the fragment into pEGFP-N1 by Red-mediated recombination ^11^. pEGFP-N1 (Takara Bio USA, Inc., CA, USA) was linearized with homologous arms by inverse PCR using primers 1 and 2 listed in Supplementary Table 1, and the PCR product was electroporated into *Escherichia coli* strain GS1783 ^11^ competent cells containing pYEbac102. After a 1-h incubation in 1 ml of LB broth, the transformants were selected on LB plates supplemented with 50 mg/ml of kanamycin at 32°C for 24 h. Single colonies were streaked onto LB (kanamycin) plates with a sterile toothpick and the plates were incubated at 32°C for 24 h. Plasmids were isolated using the QIAprep Spin Miniprep Kit (Qiagen, Hilden, Germany). The inserts were confirmed by restriction digestion of the purified plasmids. To construct pBS-Puro2ABFP-WPRE, Puro2ABFP-WPRE was amplified from pKLV2-U6gRNA5(BbsI)-PGKpuro2ABFP-W (Addgene plasmid #67974; http://n2t.net/addgene:67974; RRID:Addgene_67974) ^12^ by PCR using primers 3 and 4 listed in Supplementary Table 1 and cloned into the *Bam*HI-*Kpn*I site of pBluescript II KS (+) (Stratagene, CA, USA) using ligation mix (Takara Bio USA). To construct pBS-Puro2ABFP-WPRE-AphI, the kanamycin-resistance gene AphAI flanked by an I-*Sce*I restriction site (I-*Sce*I-*Aph*AI fragment) was amplified using primers 5 and 6 listed in Supplementary Table 1 from pEPkan-S (Addgene plasmid #41017; http://n2t.net/addgene:41017; RRID:Addgene_41017) ^13^ and inserted into the *Xho*I site of pBS-Puro2ABFP-WPRE using GeneArt Gibson Assembly EX Master Mix (Thermo Fisher Scientific). To construct pmRFP, mRFP was amplified from pDEST-12.5’RFP ^14^ by PCR using primers 7 and 8 listed in Supplementary Table 1 and cloned into the *Nhe*I-*Kpn*I site of pEGFP-C2 (Addgene, #6083-1) using the In-Fusion® HD Cloning Kit (Takara Bio USA). To construct mRFP-AphAI, AphAI was amplified using primers 9 and 10 listed in Supplementary Table 1 from pEPkan-S and inserted into the *Pst*I site of pmRFP using the In-Fusion® HD Cloning Kit (Takara Bio USA).

pICP0-*Hin*dIII was constructed by retrieving the fragment that contains the ICP0 promoter, polyA, and its coding region (GenBank: GU734771.1 map position 1232–5881) from pYEbac102 and inserting the fragment into pEGFP-C2 by Red-mediated recombination, as described above, using primers 11 and 12 listed in Supplementary Table 1. The plasmid vector encoding the MCS and the EGFP-expressing cassette was constructed by VectorBuilder (vector ID: VB221026-1563hjy, vectorbuilder.com), linearized by inverse PCR using primers 13 and 14 listed in Supplementary Table 1, assembled with the ICP0-*Hin*dIII fragment digested from pICP0-*Hin*dIII using the In-Fusion® HD Cloning Kit (Takara Bio USA), and cloned into pVB-ICP0-MCS-GFP. The fragment containing the HSV-1 packaging sequence (*pac*) (GenBank: GU734771.1 map positions 151,376–152151, 1–428) ^15^ was obtained by retrieving the fragment using primers 15 and 16, and the fragment containing the HSV-1 replication origin (*oriS*) (GenBank: GU734771.1 map position 131,461–132,421) ^16^ was obtained by retrieving the fragment using primers 17 and 18 listed in Supplementary Table 1, as described above. To construct pA-ICP0-MCS-GFP, the *pac* fragment obtained from pEGFP-C2-pac by *Hin*dIII-*Bam*HI digestion and the *oriS* fragment obtained from pEGFP-C2-oriS by *Bam*HI-*Cla*I digestion were inserted into the *Hin*dIII-*Cla*I site of pVB-ICP0-GFP using ligation mix (Takara Bio USA).

### HSV-BAC recombineering

HSVBAC/Δ27/ΔTR/ICP0n212/Δpac/EBNA1-oriP-Puro2ARFP was generated by a two-step Red-mediated mutagenesis procedure in *E. coli* strain GS1783 ^11^. pEF1a promoter was amplified from VB221026-1563hjy by PCR using primers 19 and 20 listed in Supplementary Table 1. The *Nhe*I-*Spe*I fragment, including EBNA-1 and mini-oriP, was obtained using digestion enzymes from pEB6CAG ^7^. The fragment containing the hPGK promoter and Puro2ABFP-WPRE-AphI was amplified from pBS-Puro2ABFP-WPRE-AphI by PCR using primers 21 and 22 listed in Supplementary Table 1. To construct HSVBAC/Δ27/EBNA1-oriP-Puro2ABFP, these three fragments were assembled into a linear fragment using GeneArt Gibson Assembly EX Master Mix (Thermo Fisher Scientific), the fragment was electroporated into GS1783 competent cells and recombined with the ICP27 gene of pYEbac102, followed by I-*Sce*I-enhanced deletion of the AphAI gene in GS1783. To replace the blue fluorescence protein (BFP) encoded in HSVBAC/Δ27/EBNA1-OriP-Puro2ABFP with mRFP, the mRFP-AphAI fragment was amplified by PCR using primers 23 and 24 listed in Supplementary Table 1, electroporated into GS1783 competent cells and recombined with BFP of pYEbac102, followed by I-*Sce*I-enhanced deletion of the AphAI gene in GS1783. To delete the terminal region, pEPkan-S was amplified by PCR using primers 25 and 26 listed in Supplementary Table 1, electroporated into GS1783 competent cells and recombined with the terminal region, followed by I-*Sce*I-enhanced deletion of the AphAI gene in GS1783. To insert a nonsense mutation into the ICP0 gene ^17^, pEPkan-S was amplified by PCR using primers 27 and 28 listed in Supplementary Table 1, electroporated into GS1783 competent cells, followed by I-*Sce*I-enhanced deletion of the AphAI gene in GS1783. Finally, to delete the packaging sequence, pEM7 -zeo was amplified by PCR using primers 29 and 30 listed in Supplementary Table 1 and electroporated into GS1783 competent cells.

### Establishment of 293/HSV cells

Flp-In^™^ T-REx^™^ 293 cells were transfected with HSVBAC/Δ27/ΔTR/ΔICP0/Δpac/EBNA1-oriP-Puro2ARFP, selected with 1 µg/ml puromycin, cloned from a single colony, and the resulting cells were designated 293/HSV cells.

### Preparation of HSV amplicon vectors

For Fig. 1d, 1e, Supplementary Fig. 1a, 1b, and 1d, 293/HSV cells were seeded at 3.4 × 10^5^ cells per well in a 24-well plate coated with 0.01% collagen (IFP9660, Research Institute for the Functional Peptides Co., Japan) for 10 min, washed with phosphate-buffered saline (PBS, Wako), and mock-transfected or co-transfected with pA-ICP0-MCS-GFP (400 ng) and pICP27 (100 ng) using PEI Max^TM^ (24765-100, Polysciences, Inc., PA, USA) following the manufacturer’s protocol. The conditioned medium was replaced with DMEM containing 10% FBS at 6 h and 48 h post-transfection. The transfected cells were analyzed by fluorescence microscopy and immunoblotting at 72 h post-transfection. The transfected cells and conditioned medium were freeze-thawed three times, cell debris was removed by centrifuging at 10,000 × g for 5 min twice, and the supernatants were collected. Vero cells were treated with the supernatants for 18 h and analyzed by fluorescence microscopy and flow cytometry. For Fig. 1f and Table 1, 293/HSV cells were seeded at 1.1 × 10^7^ cells per well in a 10-cm dish coated with 0.01% collagen for 10 min, washed with PBS, and then mock-transfected or co-transfected with pA-ICP0-MCS-GFP (11.2 μg) and pICP27 (800 ng) using PEI Max^TM^ (24765-100, Polysciences Inc., PA, USA), Lipofectamine^TM^ 2000 (11668019, Thermo Fisher Scientific), or Lipofectamine^TM^ 3000 (L3000015, Thermo Fisher Scientific) following the manufacturer’s protocols. The conditioned medium was replaced with DMEM containing 10% FBS at 6 h and 48 h post-transfection. The transfected cells were analyzed by fluorescence microscopy at 72 h post-transfection. The transfected cells and conditioned medium were collected and centrifuged at 10,000 × g for 5 min. The supernatant was removed and the transfected cells were resuspended in 1 ml of DMEM containing 10% FBS. The suspended cells were sonicated three times for 30 s at a strength of 8 with 20-s intervals on ice using a handy ultrasonic disruptor UR-21P (TOMY, Tokyo, Japan) and were stored at □80°C. The cell debris was removed by centrifugation at 10,000 × g for 5 min and the supernatant was filtered through a 0.45-μm syringe-tip filter attached to a 1 ml disposable syringe into a 1.5 ml collection tube, and the filtrated sample was stored at □80°C as vector stock.

### Immunoblotting

The transfected cells were washed with PBS and lysed with sodium dodecyl sulfate (SDS) sample buffer (62.5 mM Tris-HCl [pH 6.8], 20% glycerol, 2% SDS, 5% 2-mercaptoethanol) at 72 h post-transfection. The cell lysates were subjected to electrophoresis in denaturing gels and transferred to polyvinylidene difluoride membranes. The membranes were blocked with 5% skim milk in Tris-buffered saline with Tween 20 for 30 min and reacted with the indicated antibodies overnight at 4°C. The membranes were then reacted with secondary antibodies conjugated with peroxidase (GE Healthcare Bio-Sciences, IL, USA) for at least 1 h at room temperature. Bands were visualized using ECL (ImmunoStar LD, Wako) with FUSION SOLO S (Vilber, Marne-la-Vallée, France).

### Fluorescence microscopy

Fluorescent images of transfected cells were obtained at 72 h post-transfection with an OLYMPUS fluorescence microscope IX71, DP74 (OLYMPUS, Japan) at 5× magnification.

### Flow cytometry

Cells were separated from the culture plate by trypsinization, washed with PBS, centrifuged at 200 × g for 5 min, fixed with 2% paraformaldehyde solution, centrifuged at 200 × g for 5 min, and resuspended in 500 µl of PBS with 10% FBS. The fluorescence of EGFP was analyzed directly with a cell sorter SH800S (SONY, Japan). We assessed the rates of EGFP-positive cells by two-dimensional plot analysis to distinguish the EGFP-positive cells from the EGFP-negative (mock-infected) cells.

### Statistical analysis

For comparisons of two groups, statistical analysis was performed using an unpaired Student’s *t*-test. Tukey’s test was used for comparisons among multiple groups. A p-value > 0.05 was considered not significant (n.s.). All statistical analyses were performed using GraphPad Prism 7 (GraphPad Software, San Diego, CA, USA).

## Data availability

The data that support the findings of this study are available from the corresponding author upon reasonable request.

