## Supplemental Figures and Table 1 for "Development of the “EASY-HSV (Efficient And Simple high-Yield Herpes Simplex virus Vector) system” based on HSV-1 genome-maintaining HEK293 cells"

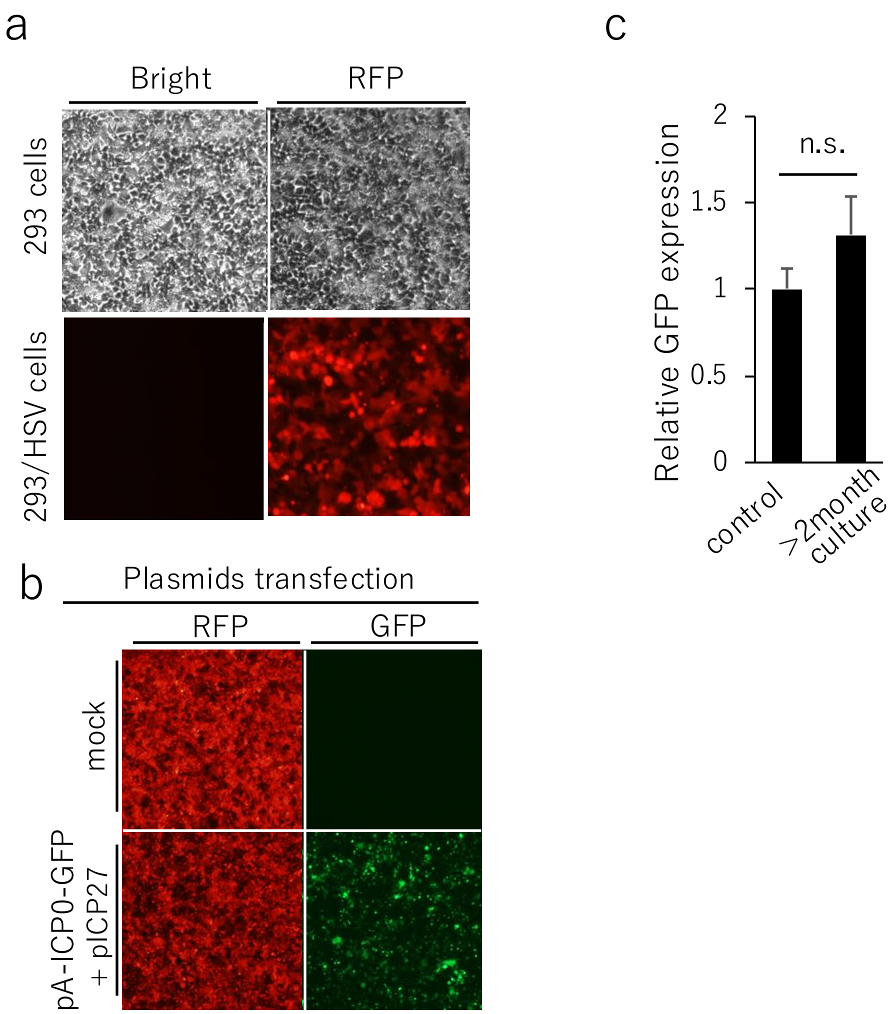


**Supplementary Figure 1: Characterization of 293/HSV cells.** (a) 293 cells and 293/HSV cells were analyzed by fluorescence microscopy. (b) Fluorescence microscopy images showing 293/HSV cells mock-transfected or transfected with pA-ICP0-GFP and pICP27. (c) HSV amplicon vector stocks were prepared using cells (control) and cells cultured for more than two months longer than the control (>2month culture). Vero cells were infected with HSV amplicon vector stocks prepared with cell lines for 18h. Data are shown as the means ± standard error of the results of 3 independent experiments and are expressed relative to the mean determined for control, which is normalized to 1. Statistical analysis was performed by Student's t-test.

### Supplementary Table 1: Primer list

| No. | sequence |
|---|---|
| 1 | tacaaggacacccaccacacccctccgctgattaccgagctctagaaggcgtaaattgtaagcgtt |
| 2 | ggacggccaattgggacccatgggcggggtcgttgggatcgtcgaccggaactccatatatgggct |
| 3 | gcggatccaattctaccgggta |
| 4 | gcgtacctgcggggaggcggcc |
| 5 | ccatctagatctcgagcagctgaagcttaccataggatgacgacgataagtag |
| 6 | gcttcagctgctcgagatctagatggatggatgcaaccaattaaccaattctgattag |
| 7 | cgtcagatccgctagcatggcctcctccgaggacgt |
| 8 | ccgggcccgcggtaccggcgccggtggagtggcggc |
| 9 | ggactcctccctgcaggacggcgagttcatctaaggatgacgacgataagtag |
| 10 | actcgccgtcctgcagggaggagtcctgggtcacaaccaattaaccaattctgattag |
| 11 | tgccatgttgggcaggctctggtgttaaccacagagccgcggcccgggctaagcttagatctcgagctcaagcttc |
| 12 | gagctcccgggagctccgcggaagacccaggccgcctcgggtgtaacgttaagcttcggaactccatatatgggct |
| 13 | ggcccgggctaagcttatcgatgctagcgg |
| 14 | cctcgggtgtaagctcgtgatacgcctatttttat |
| 15 | cccgggctgccacaggtgtaacaacaccaacagaacaccaacagcacggcgaattcaccggtcgccaccatggtga |
| 16 | cgcggccgagacgagcgagttagacaggcaagcactactcgcctctgcacggatccaagcttgagctcgagatctg |
| 17 | ggtctctccggcgcacataaaggcccggcgcgaccgacgcccgcagacggaagcttaccggtcgccaccatggtga |
| 18 | acggccaacgggcgcgcggggctcgtatctcattaccgccgaaccgggaaggatccgagctcgagatctgagtccg |
| 19 | ccccggcaccacgggtataaggacatccaccacccggccgggctccggtgcccgtcagtg |
| 20 | catcctcaagatttgcgtcctgagcctcaagccaggctagtcacgacacctgaaatggaa |
| 21 | ttagttcgtccggcggcgggatctcgacattgattattgagggtaggggaggcgcttttc |
| 22 | cgtggggcgatttgtttgaaatgttttgtttttattgtacgtacctgcggggaggcggcc |
| 23 | tctcctaacatgcggtgacgtggaggagaatcctggcccaatggcctcctccgaggacgt |
| 24 | tctttcacaaattttgtaatccagaggttgagcggccgctttaggcgccggtggagtggc |
| 25 | aggtgtccgccatttccagggcccacgacatgctcccccggtatataaaccaacgaaaagaggatgacgacgataagtag |
| 26 | cccgtatccccgttcccgcgcttttcgttggtttatataccgggggagcatgtcgtgggccaaccaattaaccaattctgattag |
| 27 | gactttatctggacgggcaatcagcggttcgccccgcggtctagactagtctagacctgaccctgggaggatgacgacgataagtag |
| 28 | ggcgacagggccctcaccgtgtgcccccccagggtcaggtctagactagtctagaccgcggggcgaacaaccaattaaccaattctgattag |
| 29 | ggggtggtagagtttgacaggcaagcatgtgcgtgcagagacgcggcctttttacggttc |
| 30 | cctgatttagttttgcagtagcgttgtttatttcgaggggataccgcacagatgcgtaag |
